# Proximal recolonization by self-renewing microglia reestablishes microglial homeostasis in the adult mouse brain

**DOI:** 10.1101/378547

**Authors:** Lihong Zhan, Grietje Krabbe, Fei Du, Ian Jones, Meredith C. Reichert, Maria Telpoukhovskaia, Lay Kodama, Chao Wang, Seo-hyun Cho, Faten Sayed, Yaqiao Li, David Le, Yungui Zhou, Yin Shen, Brian West, Li Gan

**Affiliations:** Gladstone Institutes of Neurological Diseases, Department of Neurology; San Francisco, USA; Institute for Human Genetics and Department of Neurology, University of California-San Francisco, San Francisco, USA; Department of Geography, University of Wisconsin-Madison, Madison, USA; Plexxikon Inc., Berkeley, California, USA

## Abstract

Microglia are resident immune cells that play critical roles in maintaining normal physiology of the central nervous system. Remarkably, microglia have intrinsic capacity to replenish after being acutely ablated. However, the underlying mechanisms that drive such microglial restoration remain elusive. Here, we removed microglia via CSF1R inhibitor PLX5622 and characterized repopulation both spatially and temporally. We also investigated the cellular origin of repopulated microglia and report that microglia are replenished via self-renewal, with little contribution from non-microglial lineage progenitors, including nestin+ progenitors and the circulating myeloid population. Interestingly, spatial analyses with multi-color labeling reveal that newborn microglia recolonize the parenchyma by forming distinctive clusters that maintain stable territorial boundaries over time, indicating proximal expansion nature of adult microgliogenesis and stability of microglia tiling. Temporal transcriptome profiling from newborn microglia at different repopulation stages revealed that the adult newborn microglia gradually regain steady-state maturity from an immature state that is reminiscent of neonatal stage and follow a series of maturation programs that include NF-κB activation, interferon immune activation and apoptosis, etc. Importantly, we show that the restoration of microglial homeostatic density requires NF-κB signaling as well as apoptotic egress of excessive cells. In summary, our study reports key events that take place from microgliogenesis to homeostasis re-establishment.

## Introduction

Microglia are resident parenchymal macrophages in the central nervous system (CNS). Along with perivascular, meningeal, and choroid plexus macrophages, these cells govern the innate immunity of the CNS [1]. Apart from their primary function in immune surveillance, microglia also perform a multitude of essential CNS tasks: supplying neurotrophic factors, pruning unwanted synapses, and promoting programmed cell death [2]. Consequently, microglia have often been implicated in neurodegenerative and neuropsychiatric diseases [2, 3]. In particular, loss of microglia homeostatic control, in the form of microgliosis, is considered one of the key pathological hallmarks.

Mechanistic insights into microglial homeostasis regulation have been challenging. Disease or ageing models are very informative in elucidating microgliosis initiation, but can be limiting when applied to studies of microglial homeostatic restoration because of the persistent proinflammatory signals inherent to these models. Similarly, investigating microglia homeostasis at steady-state is also difficult because of the low turn-over rate: less than 0.05–1% of microglia are proliferative in the rodent CNS [4, 5], making transcriptomic profiling of the newborn microglia technically restrictive. In contrast, microglial ablation/repopulation approach offers possibilities to follow bulk microglial population for longitudinal investigation. Roughly 90-99% of the microglial population can be reconstructed from regenerated source [6–8]. This approach is also advantageous in investigating the microglial homeostatic restoration processes given the absence of exogenous proinflammatory stimuli as often seen in disease or ageing models. Thus, examining the restoration phase represents new access to the intricate cellular and molecular principles underlying microglial homeostasis regulation.

A number of previous studies using microglial ablation approaches had been focused on the cellular origin of the regenerated microglia. A hidden CNS nestin+ progenitor pool was initially proposed, arguing lineage crosstalk with neurogenesis [6]. This theory has since been re-evaluated by several studies. Bruttger and colleagues showed that microglia can be regenerated almost entirely from self-renewal [7] and was further confirmed recently by Huang *et al* [8]. In addition, nestin+ progenitors transitioning into adult microglia could not be detected under steady-state proliferation [4]. Despite these efforts in resolving the origin debate, the molecular and cellular underpinning of how repopulated microglia return to steady-state homeostasis is poorly defined. Here, in addition to address the origin controversy, we also examined how microglia restore homeostasis in spatial redistribution, maturity, and cell density.

Microglia are territorial sentinels in the CNS. Resting-state microglia evenly tile the parenchyma, forming an intricate cellular grid with microglial processes constantly surveilling local territories [9]. This tiling pattern is largely determined during seeding of the primitive yolk sac progenitors at embryonic stage [10]. Despite adult microglia have the capacity to migrate over long distances, especially in response to neuronal injuries [9], it is hard to determine whether maintenance of microglia tiling is static or dynamic because microglia appear homogenous under conventional visualization methods. Here, we combined mosaic labeling using the “Microfetti” method that features distinctive fluorescent labels in microglia [11] and the ablation/repopulation approach to recreate spatial tiling in the adulthood and investigated microglia tiling maintenance over time. By applying spatial analyses such as Nearest Neighbor Distance (NND) and Ripley’s K-function [12], we were able to capture the spatial characteristics of microglial tiles up to 1 month.

Becoming mature microglia is a dynamic process. First, erythro-myeloid precursor cells must leave the blood island and travel from the yolk sac to colonize the developing fetal brain [10, 13, 14]. Next, microglia continue to develop in the immature neonatal microenvironment, following a precisely programmed path to become fully functional adult microglia [15]. Whether microglia require specialized developing environment for proper maturation is an open question, nor have the molecular underpinnings of microglial maturation during adult microgliogenesis been thoroughly defined. Here, we systematically dissected microglial transcriptome in newborn microglia from different repopulation stages to address these questions. We found that early stage adult-born microglia are reminiscent of the neonatal counterpart and undergo a series of transcriptional changes that eventually regain steady-state maturity. Our results show that microglia can fully mature in the adult brain. We also discovered distinctive molecular programs that are associated with microglial maturation during adult microgliogenesis. In particular, we found immune signaling belonging to the NF-κB and interferon pathways were highly evaluated, accompanied by large amount of apoptosis. Conditional knock-out of the IKKβ, a critical kinase required for NF-κB activation, caused significant defects in repopulation of microglia *in vivo*. Finally, we address how microglia density is restored to steady-state level from an overproliferation state after repopulation had completed [6–8]. We performed 5-ethynyl-2’-deoxyuridine (EdU) pulse-chase labeling to record the egress kinetics of newborn microglia. We discovered that restoration of homeostatic microglial density was achieved by slow turnover of microglia, demonstrating the striking length required for microgliosis resolution under sterile environment.

## Results

### Release from CSF1 inhibition triggers proliferation of microglia to reach homeostatic density

Microglia are resident immune cells that territorially tile the entire CNS. Microglial homeostasis is tightly controlled under physiological conditions [4]. To investigate the mechanisms underlying microglia homeostatic restoration, we depleted microglia by feeding mice with a diet containing CSF1R inhibitor PLX5622 (PLX) (Figure 1a). Administration of PLX for 2 weeks reduced the number of Iba1+ microglia by 80%. Switching to a regular control diet for 4 days restored microglia density to levels near non-treated control (Figure 1b-d). Similarly, microglial repopulation was also observed following diphtheria toxin-mediated selective ablation of microglia (Supplementary Figure 1a). Microglia density was partially restored after 1 day of repopulation and even surpassed normal density after 7 days of repopulation (Supplementary Figure 1b, c). These results indicate that repopulation is an inherent property of microglia that is independent of ablation approaches.

**Figure 1.**
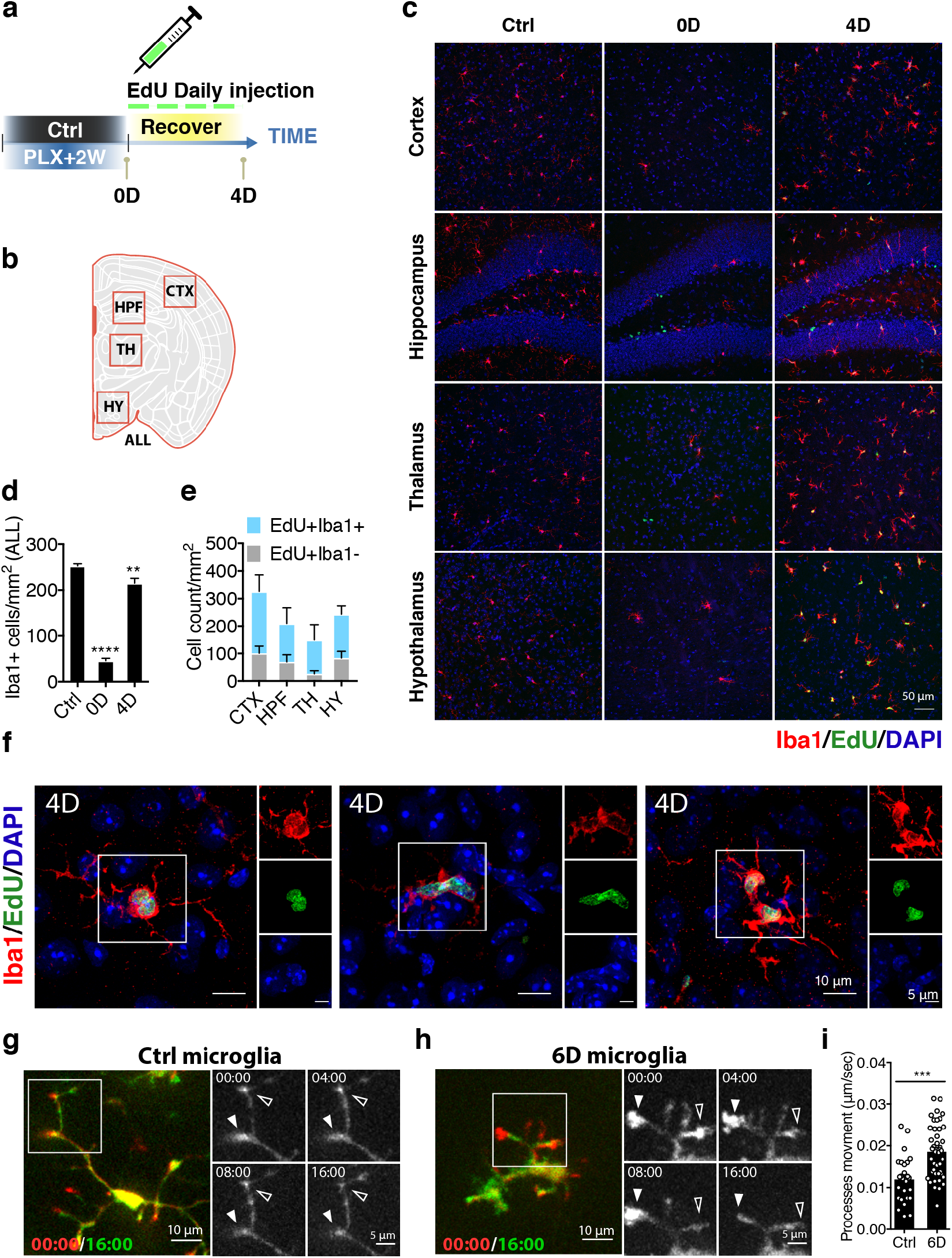
Release from CSF1 inhibition triggers proliferation of microglia to reach homeostatic density. (**a**) Schematic workflow of the experimental design. Three-month old C57BL/6 mice were treated with PLX diet then switched to normal diet for 4 days to allow microglia repopulation. Control mice were fed with normal diet for the entire duration of the experiment. EdU was administered via I.P. injection every 24 hours for 4 days. (**b**) Diagram showing selected sub-brain regions for quantification. Abbreviation: Cortex (CTX); Hippocampus (HPF); Thalamus (TH); Hypothalamus (HY); entire coronal section (ALL). (**c**) Confocal microscopy images showing microglia depletion and repopulation in different brain regions. The following markers were pseudo-colored accordingly: Iba1 (red); EdU (green); DAPI (blue). (**d**) Quantification of Iba1+ microglia density from the entire coronal section. Mean values from each animal were plotted with SEM. Ctrl (n = 5); 0D (n = 5); 4D (n = 5). One-way ANOVA with Dunnett’s multiple comparisons test was used by comparing to the control (ctrl) group. (**e**) Quantitation of proliferating microglia (EdU+ and Iba1+) and proliferating non-microglial (EdU+ and Iba1−) cells from different brain regions at day 4 of repopulation. (**f**) Confocal images showing Iba1+ (red) microglia undergoing mitosis as marked by EdU labeling. (**g, h**) Representative frames from live imaging of untreated control microglia (g) and microglia at day 6 of repopulation (h). Acute slices from CX3CR1-GFP mice were used to image microglia. A total of 16 mins were recorded. The first frame (pseudo-colored in red) is overlaid with the last frame (pseudo-colored in green). The box highlights movement of microglial processes. Extension is indicated with closed triangles while retraction is indicated with open triangles. (**i**) Quantification of the average velocity of all processes per cell in μm/sec from acute brain slices. Control (n = 3 animals, 6 slices, 26 cells); 6D (n = 2 animals, 10 slices, 43 cells). Each data point represents a single cell. Student’s t-test was applied. Statistical summary of p-value indicated as p > 0.05 (ns); p ≤ 0.05 (*); p ≤ 0.01 (**); p ≤ 0.001 (***); p ≤ 0.0001 (****).

Using EdU labeling to identify mitotic cells, we found that about 60% of repopulated microglia at day 4 were positive for EdU (Figure 1e), suggesting at least half of the repopulated microglia were derived from newborn cells. Proliferative microglia were found throughout the parenchyma, which showed pairs of proximal cells undergoing different mitotic stages (Figure 1f). Consistent with an active state where re-entry of cell cycle was engaged, repopulated microglia at day 6 showed increased motility (Figure 1g-i and Supplementary video 1a, b). Among the EdU+ mitotic population, about 10–30% did not express the microglial marker Iba1 (Figure 1e). Although the exact identity of these cells remains unclear, a small subset of the EdU+Iba1-cells express DCX+ or Olig2+ (Supplementary Figure 2c, g, f). Interestingly, nestin was found to be expressed on about 30% EdU+Iba1+ cells (Supplementary Figure 2h), but not on EdU+Iba1-cells (Supplementary Figure 2g).

### Bone marrow-derived hematopoietic myeloid cells do not contribute to the repopulated microglia pool

To directly test whether circulating monocytes would become repopulated microglia after PLX-mediated depletion, we performed bone marrow transplantation using wild-type C57BL/6J mice as recipients and ACTB-eGFP as donor mice. Circulating monocytes in the chimera mice express eGFP (Supplementary Figure 3a). To protect the blood-brain barrier from irradiation damage, a custom-fitted lead helmet was applied to the mice [16] (Supplementary Figure 3b). The chimeric mice were subjected to microglia depletion on PLX diet for 2 weeks and then switched to normal diet for repopulation. On average, chimeric mice showed 70% myeloid reconstitution as determined by the presence of eGFP in the CD11b+CD45+ population from blood or splenocyte suspension (Supplementary Figure 3c-e). Immunofluorescence staining of coronal brain slices showed very few GFP+ cell infiltrated the parenchyma, either after 14 days or 2 months of repopulation, and those few that did infiltrate did not express the microglial marker Iba1 (Supplementary Figure 3f, g), except for those in the choroid plexus (Supplementary Figure 3h). Our finding provides direct evidence that bone marrow-derived hematopoietic myeloid cells do not contribute to the repopulated microglial pool, consistent with previous studies [7, 8].

### Lineage tracing studies established that repopulated microglia are derived from a CX3CR1+ cell lineage

To further establish the origin of the repopulated microglia, we performed fate-mapping analysis using a combination of the STOP^flox^-RFP reporter [17] and the tamoxifen inducible myeloid specific driver CX3CR1-CreERT2 [18] to genetically label adult microglia (Figure 2a). This genetic labeling approach ensures that any cells originating from RFP+ cells retain RFP expression. Intraperitoneal injection of tamoxifen at a daily dose of 2 mg/mouse for 10 consecutive days resulted RFP labeling in over 85% of microglia (Figure 2b, c), a rate comparable with a previous study [18]. After microglia depletion with PLX treatment followed by repopulation, the percentage of RFP+ cell among repopulated microglia remained similar to untreated controls at ~85% (Figure 2b, d). Even with an additional round of depletion and repopulation, the percentage of RFP+ cells remained unchanged. Indeed, the second round of depletion did not seem to exhaust microglia repopulation capacity (Figure 2d). Rather, excessive microglial proliferation exceeding the steady-state level was observed. Thus, the repopulated microglia are almost exclusively derived from the remaining CX3CR1+ microglia, most likely via self-renewal.

**Figure 2.**
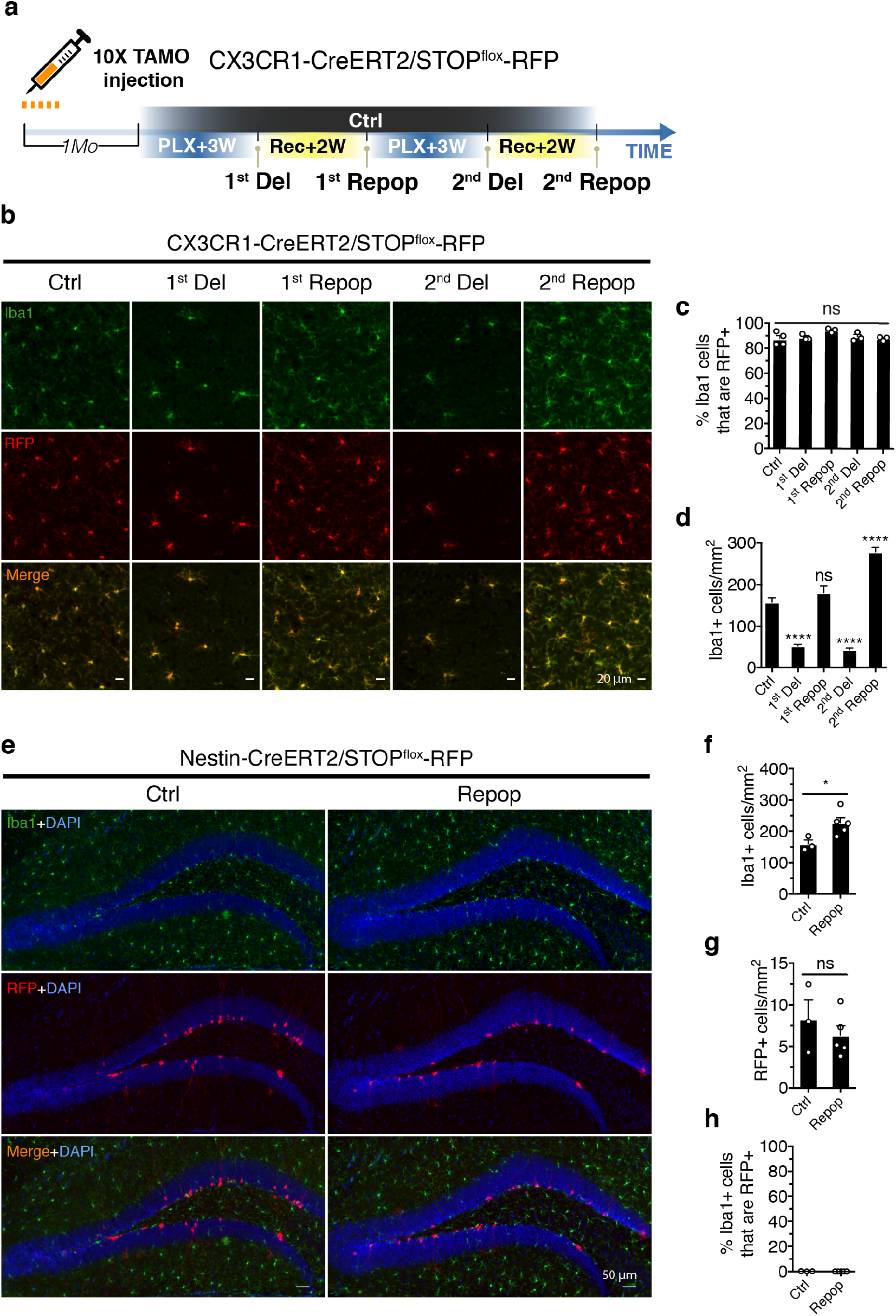
Repopulated microglia are exclusively derived from CX3CR1+ cell lineage. (**a**) Schematic workflow of the experimental design. The CX3CR1-CreERT2/STOP^flox^-RFP mice (7-9 Mo) were given 2 mg tamoxifen daily via I.P. injection for 10 days. Two weeks after the last tamoxifen administration, mice were subjected to 3 weeks of PLX treatment (1^st^ Del) before switching back to normal diet for 2 weeks (1^st^ Repop). This microglia depletion/repopulation regimen was repeated for a second round (2^nd^ Del, 2^nd^ Repop). (**b**) Representative images showing Iba1+ (green) and RFP (red) after depletion and repopulation. Images were taken from the thalamic region. (**c**) Quantification of the percentage of Iba1+ microglia that express RFP after different treatment. Mean values from each animal were plotted with SEM. One-way ANOVA was used to assess statistical differences among the groups. (**d**) Quantification of Iba1+ microglia density after different treatments. Analysis included both hippocampal and thalamic regions. Mean values from each animal were plotted with SEM. Ctrl (n = 4); 1^st^ Del (n = 3); 1^st^ Repop (n = 3); 2^nd^ Del (n = 3); 2^nd^ Repop (n = 3). One-way ANOVA with Dunnett’s multiple comparisons test was used by comparing to the control (ctrl) group. (**e**) Representative images showing nestin+ lineage (RFP+, red) and microglia (Iba1+, green) at the subgranular zone (SGZ) before and after microglia repopulation. The Nestin-CreERT2/STOP^flox^-RFP mice (4 Mo) were treated as described in (a), with a single round of microglia depletion and repopulation. (**f**) Quantification of Iba1+ microglia cell density before and after microglia repopulation. Mean values from each animal were plotted with SEM. Ctrl (n = 3); Repop (n = 4). (**g**) Quantification of cell density of nestin lineage (RFP+) before and after microglia repopulation. Mean values were plotted with SEM. (**h**) Quantification of Iba1+ microglia that express nestin lineage RFP before and after microglia repopulation. Mean values were plotted with SEM. Student’s t-test was used to compute statistical differences (f, g). Statistical summary of p-value indicated as p > 0.05 (ns); p ≤ 0.05 (*); p ≤ 0.01 (**); p ≤ 0.001 (***); p ≤ 0.0001 (****).

### Repopulated microglia are not derived from Nestin+, NG2+, and PDGFra+ progenitors

We and others have observed that, nestin, a marker of neuroprogenitor cells, appears to be transiently expressed in early-stage newborn microglia [6]’ [7]. To directly test if nestin+ progenitors can also contribute to the repopulated microglial pool, as proposed previously [6], we performed fate-mapping experiments using Nestin-CreERT2 [19] and a STOP^flox^-RFP reporter to label nestin+ progenitor cells with RFP. Neuronal precursor cells were labeled with RFP in the subgranular zone (SGZ) as expected (Figure 2e). After 3 weeks of PLX treatment followed by 2 weeks of microglia repopulation (Figure 2f), the number of RFP cells was unchanged (Figure 2g) and none of the repopulated microglia expressed RFP (Figure 2h). Thus, nestin+ progenitors do not contribute to the repopulated microglia pool. Given that a small proportion of EdU+Iba1-express Olig2+ during microglia repopulation (Supplementary Figure 2f), we tested whether oligodendrocyte precursor cells (OPCs) could give rise to microglia. STOP^flox^-RFP reporter mice were crossed with PDGFra-CreERT2 mice [20] and NG2-CreERT2 mice [21]. While PLX treatment did not affect the number of PDGFra+ progenitor cells labeled with RFP, none of the RFP+ cells became repopulated microglia (Supplementary Figure 4a–d). Similar results were seen from NG2+ progenitors (Supplementary Figure 4e-h). Combined, the data obtained from these lineage tracing experiments indicate that the repopulated microglia pool is not derived from these cell lineages but from CX3CR1+ microglia. This suggests that microglia self-renewal alone is sufficient to repopulate the entire parenchyma, as also found by Bruttger *et al*. [7] and Huang *et al* [8].

### Repopulated microglia exhibit distinctive cell clusters

To capture the spatial characteristics of repopulated microglia, we applied “Microfetti” labeling in mice featuring the Brainbow2.1 reporter driven by CX3CR1-CreERT2 to label microglia stochastically with 4 fluorescent proteins: green fluorescent protein (GFP), yellow fluorescent protein (YFP), cyan fluorescent protein (CFP), and red fluorescent protein (RFP) [11, 22]. Due to fixative induced inactivation of native fluorescent signals, we used anti-GFP antibody to detect GFP/CFP/YFP and anti-RFP antibody to detect RFP (Figure 3a). This is possible because GFP/CFP/YFP share similar protein homology but do not share any immuno-cross reactivity with RFP. After 2 weeks of PLX treatment, the CX3CR1-CreERT2/STOP^flox^-Brainbow mice were switched to a normal diet for repopulation over an extended period (up to 1 month) to examine the long-term clonal behavior of newborn microglia (Figure 3b). At a low dose of tamoxifen, uniform and sparse labeling of microglia with the Brainbow reporter was visualized via immunostaining of GFP and RFP (Figure 3c). PLX mediated depletion removed over 90% of Iba1+ microglia (Supplementary Figure 5a, b) and retained a few seeding RFP+ or GFP+ cells at day 0 (Figure 3c). After repopulation at 7 days and 1 month, clonal clusters of both RFP+ and GFP+ microglia were observed (Figure 3c). We also noticed a few scattered cells positioned between clusters that showed no clonal expansion (Figure 3c, d, f). These results suggest that first, adult microgliogenesis occurs via clonal expansion; second, residual microglia are heterogeneous in their proliferative capacity.

**Figure 3.**
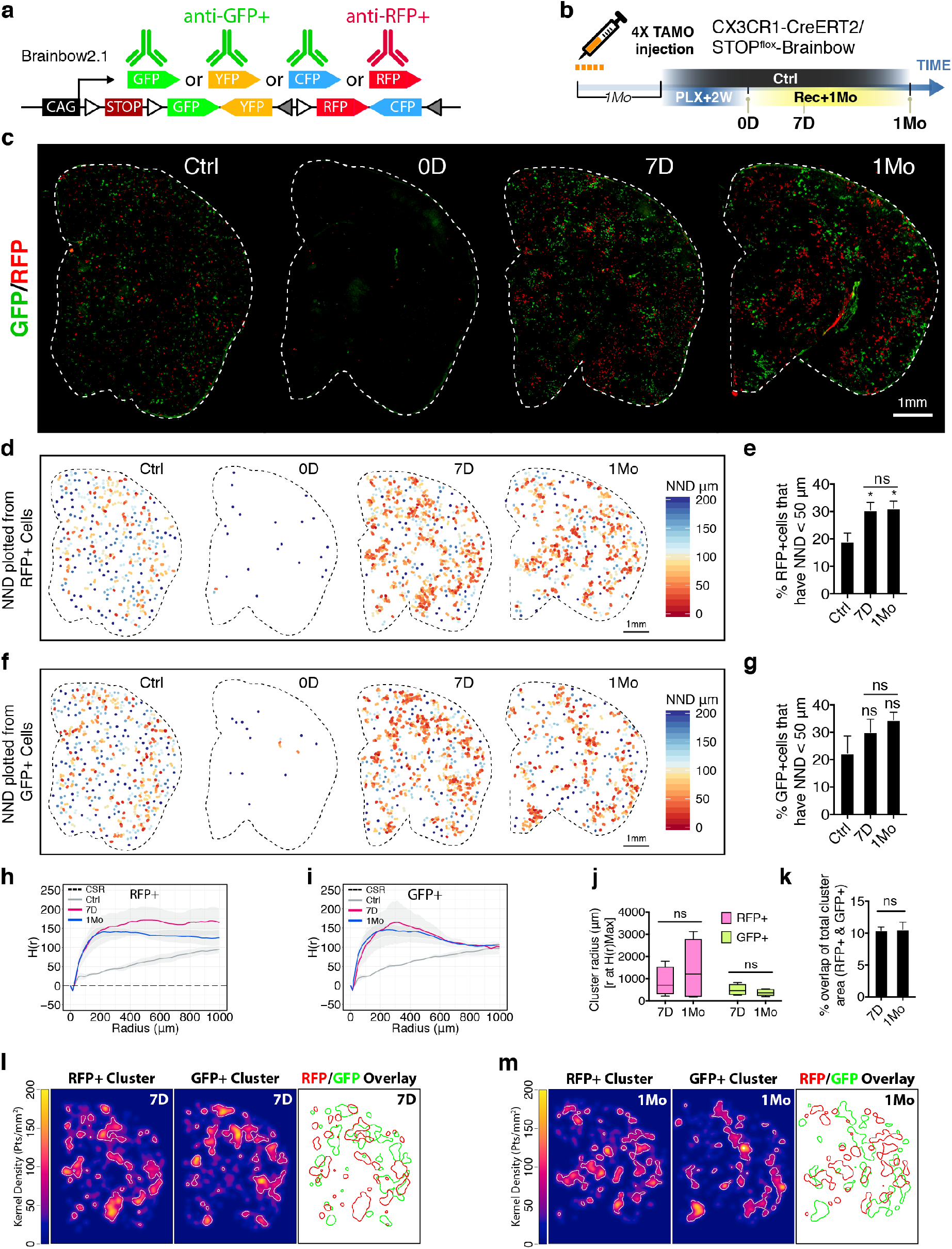
Repopulated microglia form stable clusters with minimal migratory diffusion. (**a**) Immunofluorescent staining strategy to visualize RFP+ or GFP+ microglia using the Brainbow reporter. (**b**) Experimental scheme to sparsely label microglia with Brainbow reporter. The CX3CR1-CreERT2/STOP^flox^-Brainbow mice (7-8 Mo) were given a daily dose of 2 mg tamoxifen per animal via I.P. injection for 4 days. One month after the last tamoxifen administration, mice were subjected to 2 weeks of PLX treatment before switching to normal diet for 7 days (7D) or 1 month (1Mo). (**c**) Representative images from coronal sections of RFP+ cells (red) and GFP+ cells (green). Outline of parenchyma is designated with a white dotted line. (**d**) Spatial heatmap of nearest neighbor distance (NND) reconstructed from RFP+ cells. Each dot represents a single cell which was color-coded with NND score. (**e**) Quantification of the percentage of RFP+ cells that have equal or less than 50 μm NND. Mean value from each animal was plotted with SEM. Ctrl (n = 5); 7D (n = 4); 1Mo (n = 5). One-way ANOVA with Dunnett’s multiple comparisons test was used by comparing to the control (ctrl) group. Sidak’s multiple comparisons test was used to compare 7D and 1Mo. (**f**) Spatial heatmap of nearest neighbor distance (NND) reconstructed from GFP+ cells. (**g**) Quantification of GFP+ cells as described in (**e**). (**h, i**) Plot of Ripley’s H-function analysis on RFP+ (h) and GFP+ (i) cell clustering patterns. Black dotted line represents complete spatial randomness (CSR), i.e., absence of clustering pattern. Average H(r) value from each group were plotted. Grey ribbon shades represent SEM. Ctrl (n = 5); 7D (n = 4); 1Mo (n = 5). (j) Cluster domain size estimation from H(r)Max. Box-whisker plot of cluster domain size estimation from H(r)Max (whisker: max and min; box: 25 and 75 percentile). Student’s t-test was used. (**k**) Quantification of the percentage of overlapping area of RFP+ and GFP+ clusters with respect to total RFP+ and GFP+ cluster area. The mean value from each animal was plotted with SEM. 7D (n = 4); 1Mo (n = 5). (**l, m**) 2D kernel density map showing cluster interaction between RFP+ and GFP+ cells. Representative sample from 7D (l) and 1Mo (m) were plotted. White lines mark the border of isolated cluster domains based on the top 10% of the highest kernel density. Overlay of the isolated RFP+ and GFP+ cluster contours are delineated with red and green lines, respectively. Statistical summary of p-value indicated as p > 0.05 (ns); p ≤ 0.05 (*); p ≤ 0.01 (**); p ≤ 0.001 (***); p ≤ 0.0001 (****).

### Repopulated microglia form stable clusters with minimal migratory diffusion

To study the properties of the clusters and their dynamics over time, we first applied nearest neighbor distance (NND) analysis and assigned NND values to individual cells based on their distance away from its nearest neighbor. Thus, cells inside a tightly formed cluster will have low NND values, whereas dispersed cells will have high NND values. The NND-encoded heatmap offers a global visualization of the clustering patterns and shows RFP+ cells form patches of clusters after 7 days (7D) and 1 month (1Mo) of repopulation (Figure 3d). On average, unlabeled Iba1+ microglia in control mice have an NND value of about 50 μm (Supplementary Figure 5c). Therefore, we used NND of less than 50 μm as a cut-off to identify changes in clustering patterns compared to controls. Using this criterion, there is an increase in the percentage of RFP+ cells with NND values less than 50 μm in 7D and 1Mo groups compared to the Ctrl group, suggesting spatial re-distribution in the repopulated microglia (Figure 3e). Interestingly, there was no difference in clustering pattern between 7D and 1Mo suggests that once the clusters are formed, they remain stable over time. Similar results were obtained from analyzing GFP+ cells (Figure 3f). NND analysis of GFP+ cells did not reach statistical significance (Figure 3g), most likely due to higher labeling efficiency in the GFP group since three fluorophores (CFP/YFP/GFP) were combined in this group, resulting in denser labeling in the control group.

To further confirm our results obtained from the NND analysis, we next used Ripley’s K-function analysis and its derivative H-function to survey the spatial points in a defined field with varying radii. The overall degree of “Clusterness” is estimated as a cumulative function of the radius. Theoretical complete spatial randomness (CSR) modelled by random Poisson distribution is equal to zero in H-function. A negative score in H-function indicates patterns of dispersion while a positive score indicates patterns of clustering. Consistent with our NND analysis, the H-function analysis showed that 7D and 1Mo groups shared similar clustering patterns that are quite different from control group for both RFP+ and GFP+ cell populations (Figure 3h, i). Furthermore, quantification of the radius of H-function at the maximum score [23] showed indistinguishable cluster sizes between 7D and 1Mo groups (Figure 3j), suggesting that the size of both RFP+ and GFP+ clusters remain relatively constant over time.

We next investigated whether distinct clusters could merge with neighboring clusters through cell migration and spread. Using 2D kernel density maps generated by applying kernel smoothing to actual microglia spatial points, we examined the cluster interaction between RFP+ and GFP+ cells. In order to distinguish cluster boundaries, we used top 10% density from the kernel density map as a cut-off. This method allows us to artificially outline clusters boundaries for both RFP+ and GFP+ cells at 7 days (Figure 3l) and 1 month of repopulation (Figure 3m). We then measured the relative overlapping area between RFP+ and GFP+ clusters. If a cluster merged into neighboring clusters, the overlap area would increase accordingly. The analysis showed constant overlap area from 7D to 1Mo groups (Figure 3k), suggesting that the clusters’ territories are stable with minimal migratory diffusion during the repopulation process. Together, these data suggest adult newborn microglia redistribute spatially, forming distinct clusters with patterns that are stable over the time course of repopulation.

### Early-stage newborn microglia recapitulate immature signatures of the neonatal stage

We next performed RNA-seq to profile the transcriptome of adult newborn microglia during different stages of repopulation after PLX treatment (Supplementary table 1a). Microglia were isolated from repopulated microglia of 4 days (4D), 14 days (14D) and 1 month (1Mo) after 2-weeks of PLX treatment (Figure 4a). Microglia isolated from postnatal day 4 (P4) pups were also included as a reference for the transcriptional profile of immature microglia [15, 24]. Dimensional reduction with principal component analysis (PCA) showed biological replicates from each treatment group were very closely related, as indicated by the clustering (Figure 4b). There was a clear separation of the adult microglia (Ctrl) and immature microglia (P4) clusters as expected (Figure 4b). Interestingly, early-stage newborn microglia (4D) samples clustered between the Ctrl and P4 microglia clusters, while medium-(14D) and late-stage (1Mo) microglia clustered closer with control adult microglia (Ctrl), suggesting newborn microglia undergo progressive restoration towards a steady-state microglia transcriptional profile.

**Figure 4.**
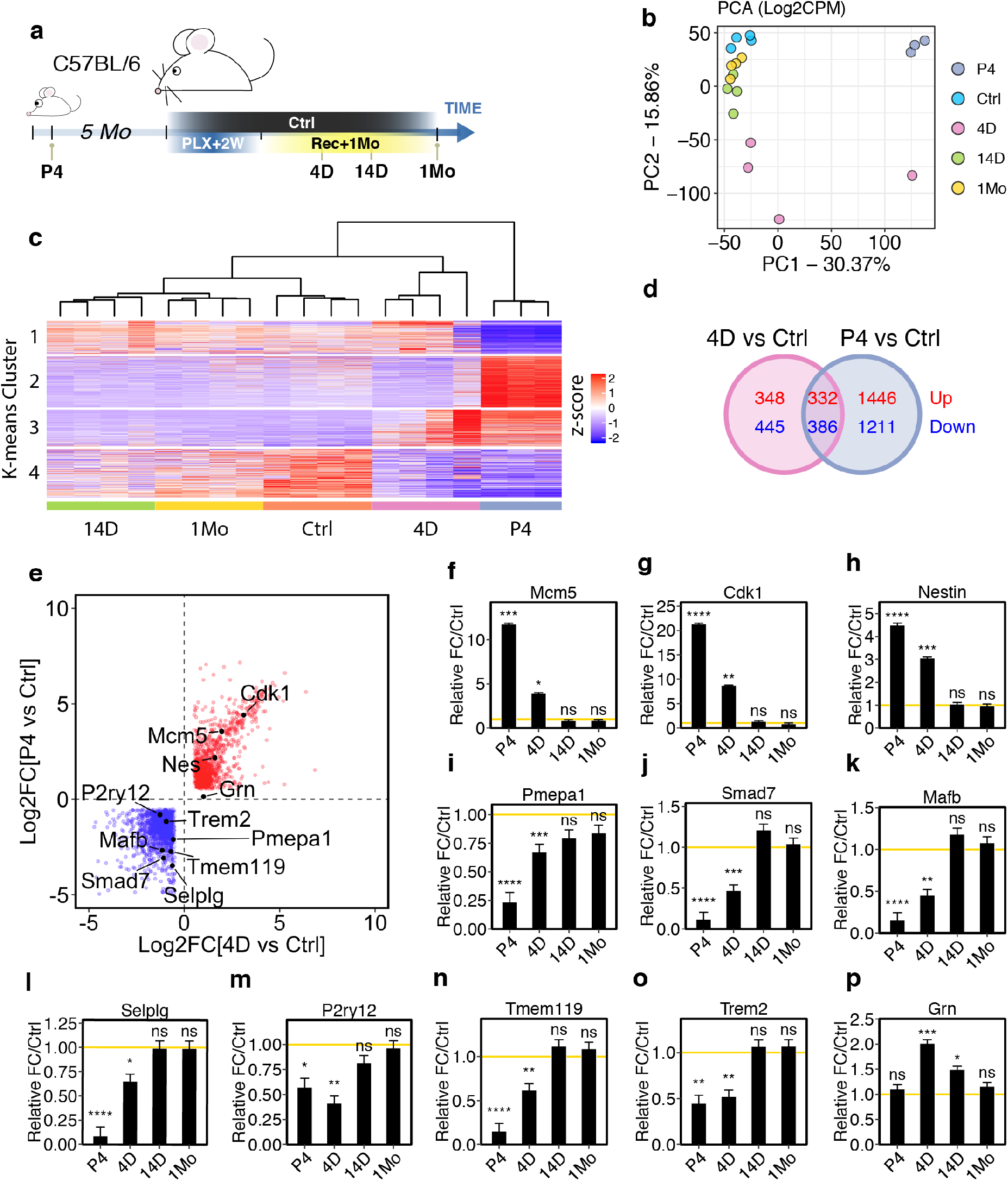
Adult newborn microglia progressively restore homeostatic maturity from a unique immature state. (**a**) Schematic workflow of RNA-seq profile experimental design. Five-month old C56BL/6 mice were treated with 2 weeks of PLX diet then switched to normal diet for 4 days (4D), 14 days (14D), or 1 month (1Mo). PCA analysis of relative gene expression variance from control (Ctrl), P4 neonatal microglia (P4), and after 4 days (4D), 14 days (14D) and 1 month (1Mo) of repopulation. The largest principle component PC1 and PC2 were used to plot the data. (**c**) Heatmap showing k-means clustering (k = 4) for relative gene expression. Dendrogram indicates hierarchical clustering of each biological replicates. Numbers of animals: Ctrl (n = 4); 4D (n = 4); 14D (n = 4); 1Mo (n = 4); P4 (n = 3). (**d**) Venn diagram comparing differentially expressed genes between 4D adult newborn microglia and P4 neonatal microglia. DE genes: Log2FC ≥ 1 or ≤ −1 and FDR < 0.05 in comparison to unperturbed adult microglia (Ctrl). Red numbers indicate upregulation while blue numbers indicate downregulation. (**e**) Scatter plot of log fold-change of differentially expresses genes of 4D microglia genes (X-axis) and P4 microglia genes (Y-axis). Red dots indicate upregulated genes and blue dots indicate downregulated genes. (**f-p**) Relative gene expression of *Mcm5* (f), *Cdk1* (g), *Nestin* (h), *Pmepa1(i), Smad7* (j), *Mafb* (k), *Selplg* (l), *P2ry12* (m), *Tmem119* (n), *Trem2* (o) and *Grn* (p). Relative fold-change was calculated in comparison to untreated control microglia. The yellow lines (y = 1) indicate normalized baseline expression of control. Statistical summary of FDR was indicated as p > 0.05 (ns); p ≤ 0.05 (*); p ≤ 0.01 (**); p ≤ 0.001 (***); p ≤ 0.0001 (****).

Next, we applied k-means clustering to arbitrarily categorize all genes into 4 distinct clusters. Among all the repopulating microglia groups, early stage newborn microglia (4D) showed the most transcriptional difference in comparison to control (Figure 4c). Interestingly, genes found in cluster 3 and cluster 4 shared the same expression directionality between 4D repopulated microglia and P4 neonatal microglia (Figure 4d). This suggests that the early stage newborn microglia might have partially reverted back to an immature developmental state. To assess the similarities between different stages of microglia, we employed a Poisson distance matrix to estimate the overall similarity between samples. The analysis showed that early stage adult newborn microglia (4D) shared 34% similarity with neonatal microglia (P4) and 55% similarity with adult microglia (Ctrl) (Supplementary Figure 6a). This suggests that 4D microglia adopted a unique immature transcriptome signature that partially overlaps with that of the neonatal microglia.

We also did analysis to specifically identify the differentially expressed (DE) genes in P4 and 4D microglia compared to control adult microglia. Roughly half of the DE genes found in 4D microglia were also present in P4 microglia (Figure 4d, e). Among all the upregulated genes shared between 4D and P4 microglia, many are involved in cell cycle regulation (Supplementary Figure 6b, c) including *Mcm5* and *Cdk1* (Figure 4f, g). Importantly, *Nestin* was also identified as an early microglia gene (Figure 4h). *Nestin* expression returned to control levels after 14 days, further validating an earlier observation that nestin was merely expressed in newborn microglia as an immature marker rather than being a *bona fide* microglial progenitor marker (Supplementary Figure 2d, h). On the other hand, among all the downregulated genes shared between 4D and P4 microglia, a large fraction of genes was involved in Mitogen Activated Protein Kinase (MAPK) pathway and Transforming growth factor β (TGF-β) signaling (Supplementary Figure 6d, e). TGF-β signaling is critically important for microglia development and homeostatic maintenance [24]. TGF-β signaling-related genes such as *Pmepa1* and *Smad7* were found to be downregulated in 4D microglia (Figure 4i, j).

### Newborn microglia re-express mature markers at later stages of repopulation

In addition, previously established mature microglia genes such as *MafB* and *Selplg* [15] were also downregulated in 4D microglia but restored after 14 days (Figure 4k, l). Furthermore, mature microglia markers such as *P2ry12* and *Tmem119* [15, 24, 25] were significantly downregulated in 4D microglia and reverted to normal levels from 14 days onwards (Figure 4m, n). These results indicate that newborn microglia regain maturity over time and was further validated in morphological analysis and immuno-staining of P2ry12 and Tmem119 protein expression in separate experiments (Supplementary Figure 7). Interestingly, disease-associated microglial genes such as *Trem2* and *Grn* were also differentially expressed in 4D microglia (Figure 4o, p). Collectively, the results gathered from the RNA-seq analyses suggest that adult microgliogenesis involves maturation steps that initiate from a unique immature state that is reminiscent of neonatal microglia.

### Adult newborn microglia exhibit distinctive temporal transcriptome profiles

To further dissect the underlying programs associated with microglia maturation, we classified four different gene sets based on their distinctive differential expression patterns during each of the repopulation stages (Figure 5a, b). The differential expression was defined as two-fold change with FDR < 0.05. We defined the first set of genes as fast-returning genes which are differentially expressed only in 4D microglia but returned to homeostatic control levels in 14D microglia (Supplementary table 1b). The second set of genes were defined as medium-returning genes which are differentially expressed in 4D and 14D microglia, but not in 1Mo microglia (Supplementary table 1c). The third set of genes were defined as slow-returning genes which are differentially expressed only in 1Mo microglia (Supplementary table 1d). In addition, we also uncovered a fourth set of genes that were only differentially expressed in 14D microglia and hence referred to as delayed-response genes (Supplementary table 1e). We applied gene set enrichment analysis (GSEA) with molecular signatures database (MSigDB) on these four gene sets [26, 27]. Comparison of the fast-returning genes with the hallmark gene dataset in MSigDB revealed enrichment of genes involved in cell division such as E2F target genes and genes involved in the G2M checkpoint or mitotic spindle (Figure 5c). Indeed, we found cell cycle-related genes such as *Ccnb2*, *Cdc20* and *Cdc25b* were highly expressed in 4D microglia but returned to homeostatic levels in 14D microglia (Figure 5 d-f). This result further supports our earlier observation that reentry of cell cycle is an early step in microglia repopulation (Figure 1f).

**Figure 5.**
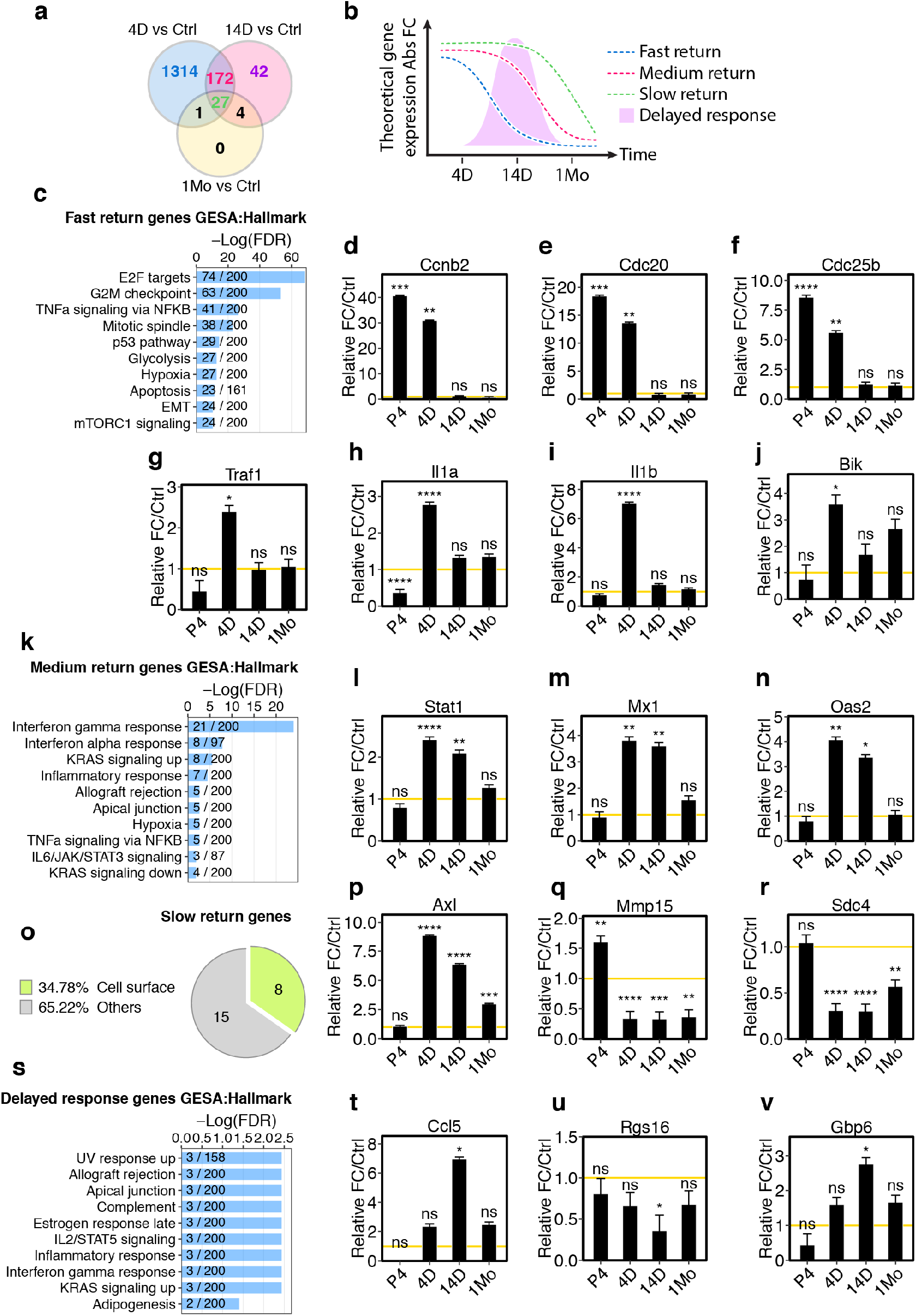
Adult newborn microglia engage stepwise program to restore steady-state gene signature. (**a**) Venn diagram comparing differentially expressed genes among 4D, 14D, and 1Mo adult newborn microglia. DE genes: Log2FC ≥ 1 or ≤ −1 and FDR < 0.05, in comparison to unperturbed adult microglia (Ctrl). (**b**) Schematic diagram illustrating gene sets with differential rates of homeostatic return. (**c**) GSEA analysis of fast return genes using the hallmark gene set database. The 10 most-enriched pathways are shown. Fraction on the bar graph indicates the total number of genes curated in the database for the specified term (denominator) and the number of overlapping genes in the enquiry RNA-seq datasets (numerator). (**d-j**) Relative gene expression of *Cdcnb2* (d), *Cdc20* (e), *Cdc25b* (f), *Traf1* (g), *Il1a* (h), *Il1b* (i) and *Bik* (j). (k) GSEA analysis on medium return genes using the hallmark gene set database. (**l-n**) Relative gene expression of *Stat1* (l), *Mx1* (m) and *Oas2* (n). (**o**) GO (cellular component) analysis on slow return genes. (**p-r**) Relative gene expression of *Axl* (p), *Mmp15* (q) and *Sdc4* (r). (s) GSEA analysis on “delayed response” genes using the hallmark gene set database. (**t-v**) Relative gene expression of *Ccl5* (t), *Rgs16* (u) and *Gbp6* (v). For gene expression profiles, relative fold-change was calculated in comparison to untreated control microglia. Yellow lines (y = 1) indicate normalized baseline expression of control. Statistical summary of FDR was indicated as p > 0.05 (ns); p ≤ 0.05 (*); p ≤ 0.01 (**); p ≤ 0.001 (***); p ≤ 0.0001 (****).

### Adult newborn microglia engage a stepwise program to restore homeostatic gene expression

Gene enrichment analyses of fast-returning gene revealed a high degree of overlap with NF-κB pathway-related genes (41 out of 200 genes) such as *Traf1*, *Il1a*, and *Il1b* (Figure 5 g-i). Both *Il1a* and *Il1b* are also associated with cell death. Interestingly, apoptosis-related genes (23 out of 161 genes) were also among the most enriched genes, including *Bik* (Figure 5j). Next, we examined the medium-return genes. Gene enrichment analysis showed strong representation of genes involved in interferon (IFN) pathways such as *Stat1*, *Mx1*, and *Oas2* (Figure 5 k-n). The enrichment of IFN genes indicates inflammatory activation during early and middle phases of microglia repopulation that was resolved after 1 month. The gene enrichment analysis for the slow-return gene set did not have statistically significant overlapping pathways. However, gene ontology analysis focused on cellular components revealed 34% of slow-return genes are cell surface related (Figure 5o). Among the most upregulated genes was *Axl* (Figure 5p), an important receptor involved in microglial phagocytic clearance of dead cells [28]. Other downregulated genes include *Mmp15* (Figure 5q), a metalloproteinase involved in extracellular matrix remodeling [29], and *Sdc4* (Figure 5r), which is an important cell surface adhesion molecule involved in cell activation and migration [30].

The delayed-response gene set revealed modest enrichment of GSEA hallmark genes (Figure 5s) with genes involved in inflammatory activation consistently represented in many categories. *Ccl5*, also known as RANTES, is an important chemotactic cytokine that modulates T cell migration [31] and was found to be highly expressed (Figure 5t). *Rgs16*, which antagonizes inflammatory activation in monocyte [32], was significantly downregulated (Figure 5u). *Gbp6*, an interferon inducible guanosine triphosphatases, was also found to be highly expressed (Figure 5v). Altogether, pathway enrichment from each gene set revealed a functional snapshot of distinct stages in microglial homeostatic restoration.

### NF-κB pathway is partially required for microglia repopulation

Among the fast-return genes, an intricate network of NF-κB pathway-associated genes was identified (Figure 6a). To understand the biological significance behind the close association of NF-κB signaling during early-phase microglia repopulation, we conditionally knocked-out kappa-B kinase subunit beta kinase (IKKβ) in microglia using CX3CR1-CreERT2/ IKKβ^F/F^ mice (Figure 6b). This approach successfully suppresses microglia activation mediated by the NF-κB pathway [33]. Next, we examined the requirement of NF-κB signaling during microglia repopulation. Experimental mice were treated for 2 weeks with PLX diet and allowed for microglial repopulation to occur over 4 days. Interestingly, IKKβ deletion showed a modest but statistically significant impairment in microglia repopulation density compared to Cre-control (Figure 6c, d). These results suggest that NF-κB signaling is at least partially required for the early phase of microglia repopulation.

**Figure 6.**
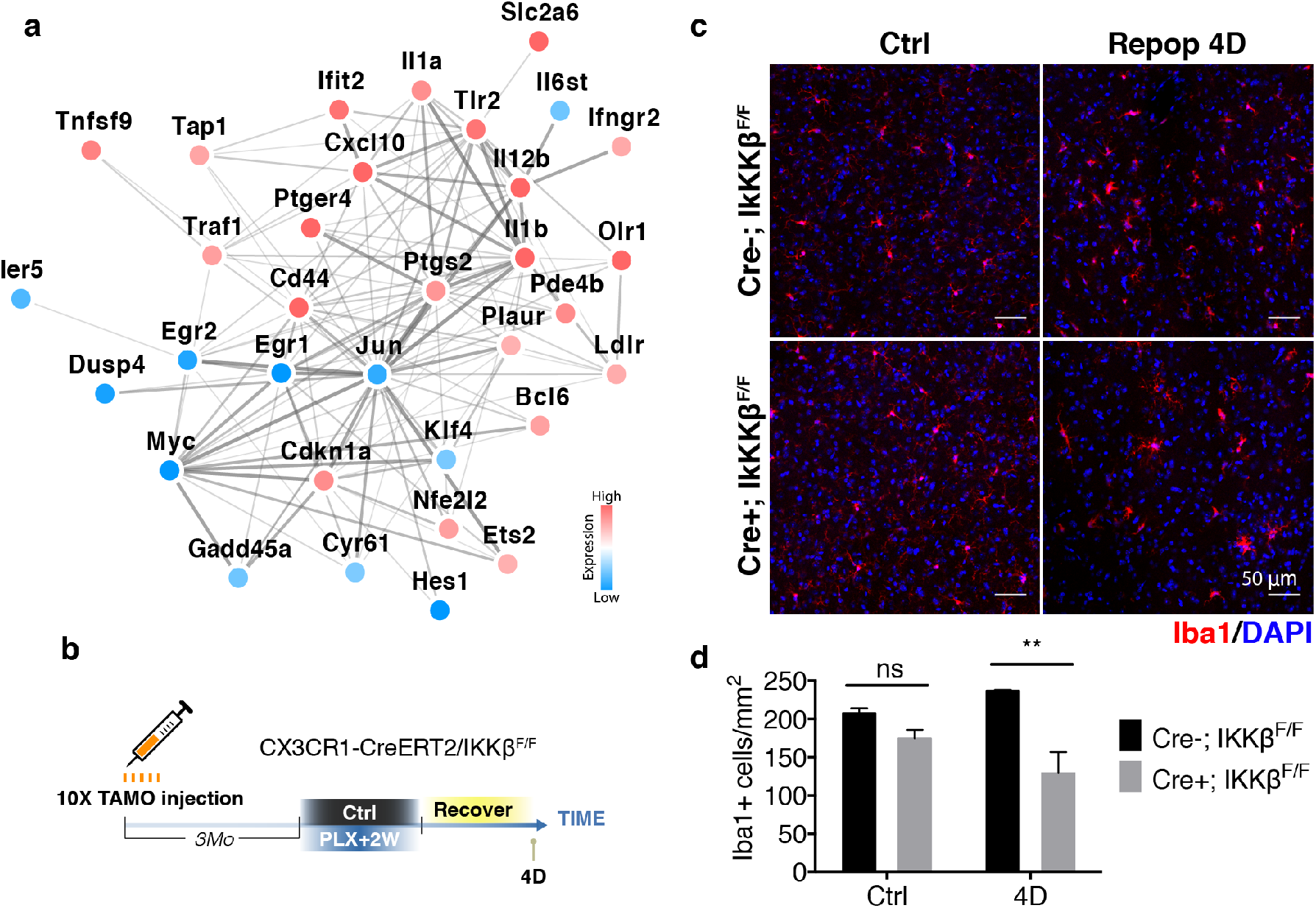
NF-kB pathway is partially required for microglia repopulation. (**a**) Network analysis using STRING database with NF-kβ signaling-associated genes identified from the fast return gene set. Relative expression of each gene is color coded as shown in the legend. (**b**) Experimental scheme for conditional knockout of IKKβ. Three months before the experiment, mice (7-8 Mo) were given a daily dose of 2 mg tamoxifen by I.P. injection for 10 days. Mice were then treated with PLX diet for 2 weeks before switching to normal diet for 4 days (4D). (**c**) Representative confocal images of control microglia and repopulated microglia (4D) with or without IKKβ deletion. Images were taken from the thalamic region. (d) Quantification of entire coronal section for control and repopulated (4D) microglia density. Mean value from each animal was plotted with SEM. Three mice (n = 3) for each group was used. One-way ANOVA with Sidak’s multiple comparisons test was used to compare IKKβ control (Cre−) and IKKβ deletion (Cre+). Statistical summary of p-value indicated as p > 0.05 (ns); p ≤ 0.05 (*); p ≤ 0.01 (**); p ≤ 0.001 (***); p ≤ 0.0001 (****).

### Cell death is associated with microglia proliferation

Apoptosis-related genes were also highly represented among fast-return genes (Figure 7a). To validate the involvement of cell death during microglia repopulation, we measured cell-death using TUNEL assay at different stages of microglia repopulation. Remarkably, the number of TUNEL-positive cells was dramatically increased in the parenchyma during the peaks of repopulation in 4D and 8D brains (Figure 7b, c). The presence of these TUNEL+ cells is not directly caused by PLX treatment since we did not observe such cell death at the end of PLX administration (0D) (Figure 7a-c). Interestingly, the TUNEL+ cells were resolved in 14D and 4Mo brains (Figure 7d), suggesting the existence of an inherent clearance mechanism. We next compared the percentage of TUNEL+ cells in mitotic microglia (EdU+Iba1+) versus non-mitotic microglia (EdU-Iba1+) at both 4D and 8D (Figure 7e, f). This comparison revealed that proliferative microglia are more likely to undergo cell death (Figure 7g). To rule out the possibility that the increased cell death might be an artifact of cell toxicity caused by EdU labeling, we compared the number of TUNEL+ cells in samples with or without EdU injection at 4D and 14D. These data suggest that EdU labeling does not skew the amount of cell death (Supplementary Figure 8a, b). Thus, active cell death is closely associated with microglia proliferation.

**Figure 7.**
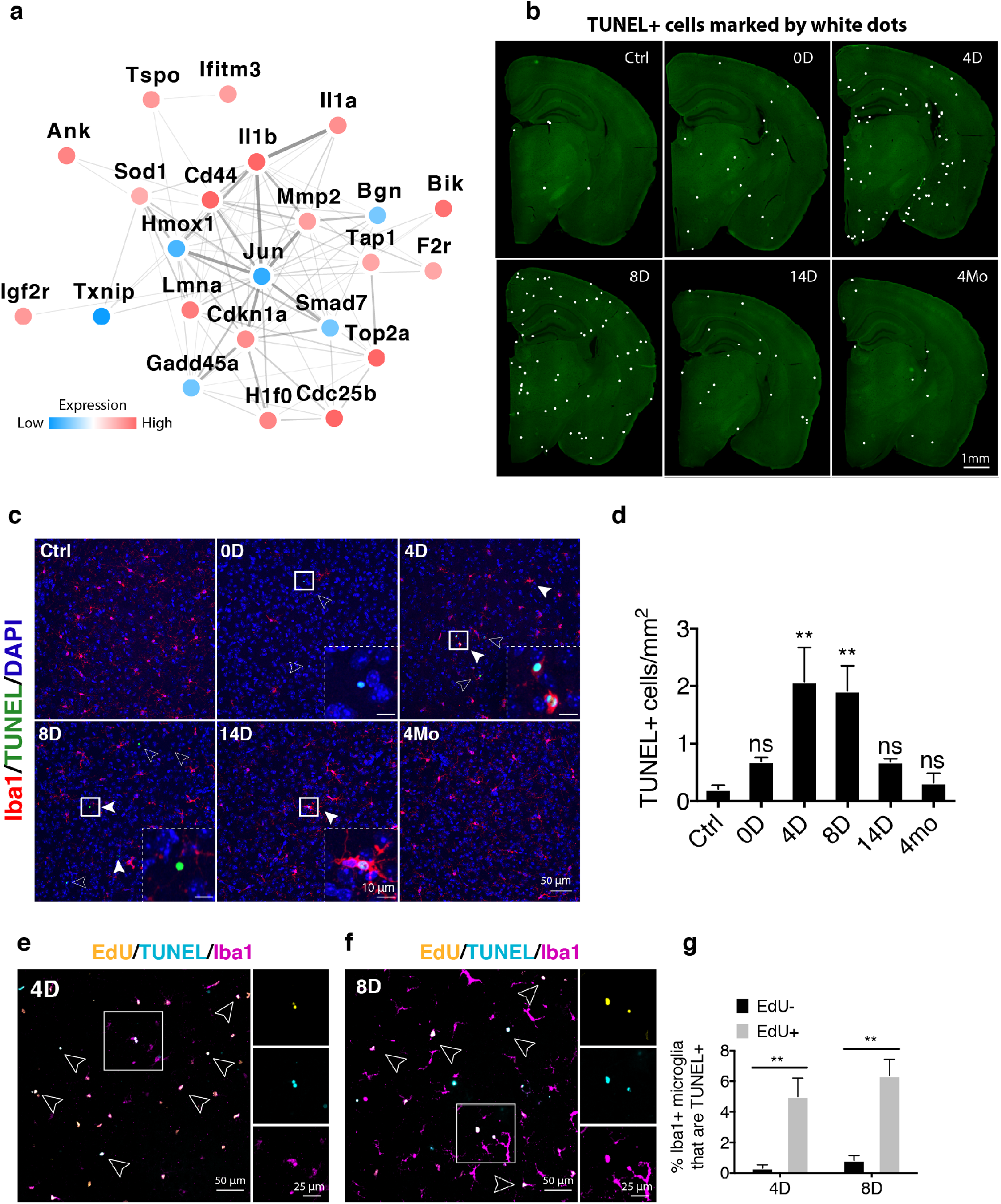
Cell death is associated with microglia proliferation. (**a**) Network analysis using the STRING database with apoptosis-associated genes identified from the fast return gene set. Relative expression of each gene is color coded as shown in the legend. (**b**) Representative images of entire coronal sections. Mice (3-5 Mo) were treated for 2 weeks with PLX diet then switched to regular diet and harvested at various time points. TUNEL+ cells are marked with white dots. (**c**) Representative confocal images of Iba1+ (red) microglia at different repopulation time point and TUNEL staining (green, highlighted with arrow heads). Images were taken from the amygdalar area. (**d**) Quantification of TUNEL+ cells density from entire coronal sections (±SEM). Ctrl (n = 5); 0D (n = 6); 4D (n = 5); 8D (n = 6); 14D (n = 7); 4Mo (n = 3). Oneway ANOVA with Dunnett’s multiple comparisons test was used by comparing to the control (ctrl) group. (**e, f**) Representative confocal images of Iba1+ (magenta) microglia that co-localize with either TUNEL (green) and/or EdU (red) at day 4 (d) and day 8 (e) of repopulation. EdU was given via I.P. injection every 24 hours during every day of repopulation after switching to normal diet. (**g**) Quantification of the percentage of TUNEL+ microglia (Iba1+) that are either EdU+ or EdU-Mean value from each animal was plotted with SEM. 4D (n = 5); 8D (n = 6). Student’s t-test was used to compute statistical differences. Statistical summary of p-value indicated as p > 0.05 (ns); p ≤ 0.05 (*); p ≤ 0.01 (**); p ≤ 0.001 (***); p ≤ 0.0001 (****).

### Microglia homeostatic density is reestablished through steady turnover

Interestingly, we and others have consistently observed that many mice developed an over-proliferative phenotype that resembles microgliosis during the first week or so of microglia repopulation (Supplementary Figure 5b) [6–8]. To determine how long it takes for repopulated CNS to restore homeostatic microglia density, we extended the recovery phase up to 4 months (Figure 8a). Removal of over-populated microglia is likely accompanied by egress of newborn cells. Thus, we applied a pulse-chase approach using EdU labeling to measure the turnover rate of newborn microglia over time by administrating daily EdU injections during the first 4 days of repopulation (Figure 8a). (Figure 8a). Time-course measurement of the amount of EdU+Iba1+ fitted with a non-linear one-phase decay model revealed steady turnover of newborn microglia with an estimated half-life of 111.3 days (Figure 8b, c). A much shorter life-span was determined for EdU+Iba1-cells (T_1/2_ = 21.44 days). The non-linear one-phase decay model assumes no additional proliferation and was confirmed by the absence of proliferation marker Ki-67 (Supplementary Figure 9). By this rate of turnover, over-proliferated newborn microglia required up to 2 months to fully egress (Figure 8d). Dead newborn microglia were removed quickly as shown earlier by the lack of TUNEL+ cells accumulation (Figure 7b, d). This is likely mediated by microglia self-engulfment as shown by examples of microglia forming phagocytic cups around dead microglia in both 14D and 4Mo brains (Figure 8e, Supplementary video 2a-d). Thus, restoration of microglia density seemed to involve steady turnover and removal of dead microglia via phagocytosis.

**Figure 8.**
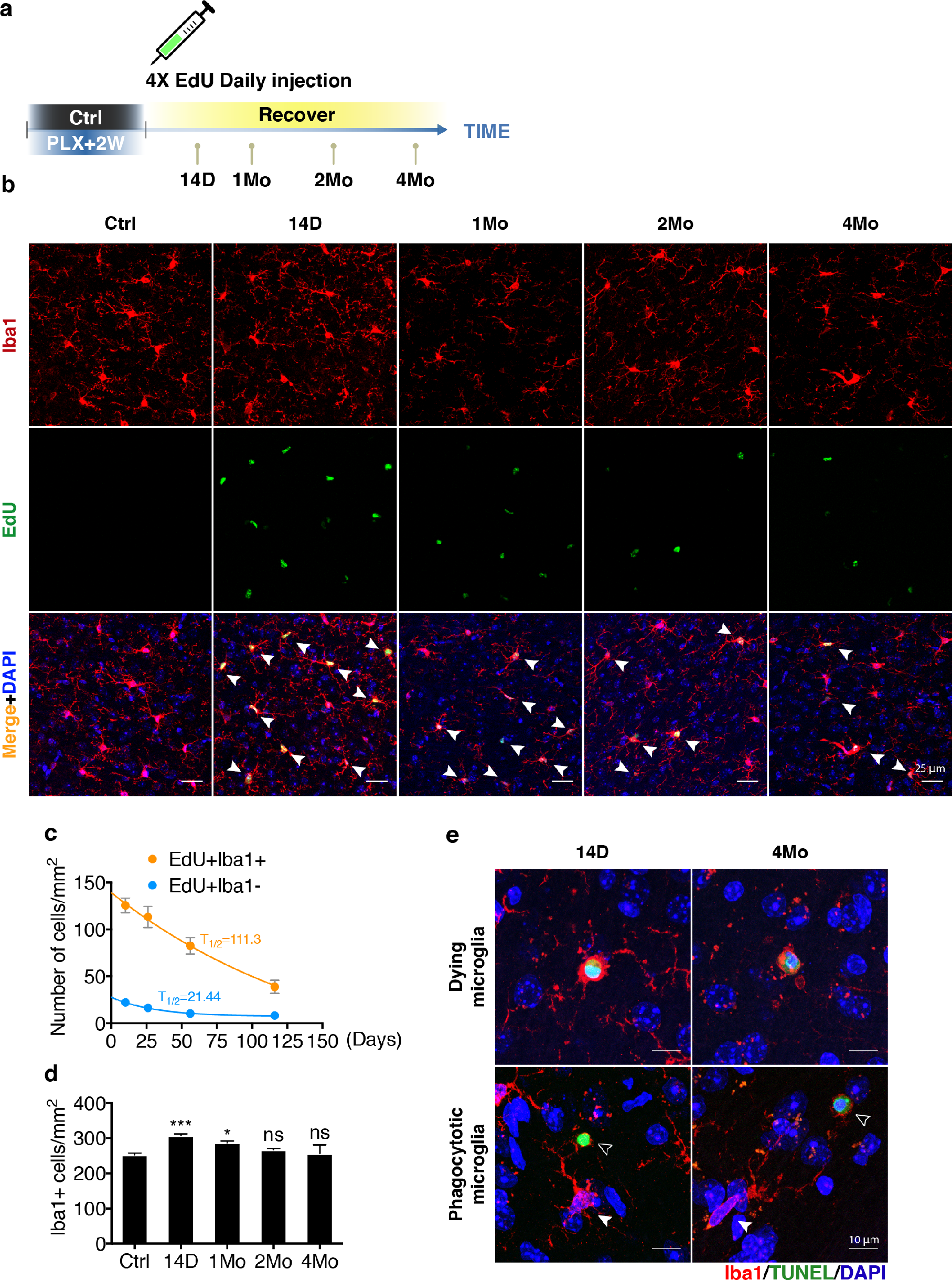
Homeostasis of adult newborn microglia is re-established through steady turnover. (**a**) Experimental scheme of pulse-chase experiments to determine newborn microglia longevity with EdU. Mice (3-5 Mo) were treated with PLX diet for 2 weeks. EdU was injected via I.P. every 24 hours during the first 4 days of repopulation. Mice were analyzed at 2 weeks (14D), 1 month (1Mo), 2 months (2Mo) and 4 months (4Mo) after switching to normal diet. (**b**) Representative confocal images of Iba1 (red) and EdU (green) at different time points. Images were taken from the cortical region. (**c**) One-phase exponential decay non-linear model fitted on EdU+Iba1+ (orange line) and EdU+Iba1- (blue line) cell density over time. T_1/2_ indicates half-life (days) of each population. Error bar indicates SEM. Animals used: 14D (n = 4); 1Mo (n = 6); 2Mo (n = 6); 4Mo (n = 3). (**d**) Quantification of total Iba1+ microglia density at different time points. Mean values from each animal were plotted with SEM. One-way ANOVA with Dunnett’s multiple comparisons test was used by comparing to the control (ctrl) group. (**e**) Representative confocal images showing TUNEL+ microglia and microglia forming a phagocytic cup around TUNEL+ cell at either 14D or 4Mo.

## Discussion

Our current study employed a chemical microglia ablation approach to query the mechanistic details of microglia homeostatic regulation. Remarkably, microglia seem to have inherent memory of their steady-state signature as repopulated microglia eventually became almost identical to resting state microglia. The data presented here piece together a series of events that take place during microglia homeostatic restoration in adulthood. First, the depleted microglia pool expands through self-renewal that partially requires NF-κB signaling. Second, microglia recolonize the parenchyma from proximal clonal expansion and maintains stable spatial clusters. Third, newborn microglia re-established maturity in a process that involves mitosis, apoptosis, interferon pathway activation, and surface molecule re-expression. Finally, homeostatic density is reached after egress of excessive newborn cells. This time course illustrates how microglia seem to have an inherent memory of their steady-state signature as repopulated microglia are eventually able to return to their resting state microglial transcriptional profile and homeostatic density.

The existence of CNS resident microglia precursors cells has been controversial [6]. In line with other studies [7, 8, 34, 35], our results demonstrate that the microglia pool can be replenished solely by self-renewal with limited contribution from either internal CNS progenitors or external circulating peripheral precursors. In the BMT chimera experiment, we observed GFP+Iba1+ in the choroid plexus (Supplementary Figure 3h). This observation is in line with recent findings that choroid plexus macrophages represent a distinctive parenchyma myeloid population that constantly receives cellular exchange with circulating monocytes [1]. Unexpectedly, we discovered an uncharacterized group of cells that are highly proliferative after release from CSF1 inhibition (Figure 1e, EdU+Iba1−). The mitotic nature of these cells was visualized by their presence of EdU labeling. Testing against a panel of common CNS markers, these cells appeared negative for Iba1, GFAP, NeuN, and S100β (Supplementary Figure 2). Although a small subset of them appeared to be DCX+ or Olig2+, the vast majority of cells indicate a non-microglial cell type that remains to be further characterized. These cells were previously considered to be microglia progenitors mainly because of their fast disappearance during repopulation, which was thought to be a result of differentiation into mature microglia cells [6]. Nonetheless, lineage-tracing experiments using CX3CR1-CreERT2 unambiguously demonstrated that microglia are repopulated solely by self-renewal.

Spatial characteristics of microgliogenesis were largely inaccessible in the past. Stochastic labeling in microglia using the “Microfetti” approach now opens new possibilities to extract spatial information during microglial tiling [11]. Using this approach, Tay *et al*. reported that microglia underwent clonal expansion in response to facial nerve injury [11]. In agreement with their finding, we found microgliogenesis under the empty niche condition also proliferate through clonal expansion. We also noticed that not all seeding cells initiated clonal expansions, as evidence by few scattered cells positioned between clusters (Figure 3c, d, f). This reflect the heterogenous nature of microglia, arguing the potential existence of sub-population of microglia with differential “stemness”. To examine the stability during microglial tiling maintenance, we followed the retiled parenchyma over the course of 1 month. We found that microglia only migrate during clonal expansion. Once the empty niche is filled, tiling boundaries remain stable up to 1 month, suggesting that once homeostasis is reached, microglia tiling is static.

Originating from yolk sac progenitors, microglia follow a series of transcriptional programs to become mature microglia [15]. Recent studies showed that adult microglia maturity is highly influenced by inflammatory signals [15, 36]. Surprisingly, exposure to an immune activing agents during pregnancy is able to shift microglia to a more mature stage [15]. On the contrary, adult microglia deprived of any immune stimuli (using a germ-free environment) displayed immature morphology, immature transcriptional signatures, and a dampened response to LPS challenge [15, 36]. These results suggest that microglia maturity is not a simple continuum of their birth age but rather their maturity is shaped by external factors such as immune activation.

In support of the necessity of inflammatory engagement during microglial maturation, we found genes associated with NF-κB signaling associated genes were highly enriched during the early phase of microglia repopulation (Figure 5c). Many of these genes are important inflammatory regulators such as Il-1a and Il-1b (Figure 5h, i). Inhibiting NF-κB pathway via conditional knockout of IKKβ resulted in partially impaired microglia repopulation (Figure 6c, d). Furthermore, we and others have found that adult microglia renewal is closely associated with cell death [4]. Although the biological significance for this cell death process awaits further clarification, we speculate these apoptotic cells might serve as “maturation trainers” for immature newborn microglia, given that apoptotic cells are highly inflammatory [37]. In support of this idea, activation of the interferon pathway occurred immediately after apoptosis (Figure 5k).

Microglia have historically been believed to be a stable cell population, a notion that [5] has been recently challenged [4]. By quantifying the microglia proliferation rate under steady-state, it was estimated that it takes 96 days to renew the entire rodent microglia pool [4]. However, a recent study by Tay *et al*. used a similar approach and concluded that microglia are extremely stable under homeostatic condition, with a turnover rate of over 41 months in cortex, 15 months in hippocampus, and 8 months in olfactory bulb [11]. These studies are advantageous in that microglia were examined in an unperturbed state. However, caveats are also present. First, assumptions on microglia cell cycle duration have to be made. Second, labeling efficiency associated with nucleotide analogs such as EdU or BrdU might not capture all proliferating microglia, thus underestimating the actual turnover rate. Third, sampling power is greatly limited since only a handful of cells can be captured at a given time. Here we implemented an EdU pulse-chase approach that directly measured the decay rate of newborn microglia, and showed that newborn microglia have a half-life of 111.3 days. (Figure 8c). In other words, adult born microglia have a lifespan of 7.5 months such that a rodent brain could renewal its entire microglial pool roughly 5 times in its lifetime. However, since our measurement started from repopulation day 14 when microglial homeostasis was not yet fully restored, 7.5 months may still be an underestimate of the actual microglia longevity. Indeed, a recent study by Füger *et al*. recorded microglia longevity via single cell imaging and reported microglia longevity is around 15 months [38]. Combined, these data suggest microglial egress can be extremely slow. This is perhaps why microgliosis is especially persistent and require months to fully resolve even in non-pathological brains (Figure 8). Therefore, strategies that can accelerate microglial turnover might be of great therapeutic value in treating neurological diseases where microgliosis is a prominent feature.

The tremendous stability associated with microglia density over the lifetime of a rodent brain is quite remarkable [4]. Even after acute ablation, microglia are able to return to the same homeostatic density. The mechanisms by which microglia are able to remember their homeostatic density is still unclear. One possible way could be due to contact inhibition. After microglia ablation, residual microglia isolated in space would lose contact inhibition exerted by neighboring microglia, thus allowing them to freely multiply. Interestingly, microglial expansion did not stop immediately after reaching homeostatic density and over-proliferation preceded the return to steady-state density (Supplementary Figure 5b) [6–8]. This suggests that the innate regulatory mechanism responsible for controlling density was switched off during microglia repopulation. Coincidentally, many of the surface molecules were still expressed at lower levels 1 month after repopulation, including Syndecan-4, which is an important regulator of contact inhibition during cancer metastasis [30]. The exact molecular machinery that sets the default homeostatic density needs to be investigated in future work.

## Acknowledgements

We want to thank Dr. David Julius at University of California San Francisco for the P2ry12 antibody; Dr. Michael Karin at University of California San Diego and Dr. Katerina Akassoglou at Gladstone Institutes for the Ikkβ^F/F^ mice; Dr. Shahzada Khan at Gladstone Institute for flow cytometry assistance; Dr. Meredith Calvert and Ms. Janna Abad at the Gladstone Histology & Light Microscopy Core for their imaging assistance; Mr. Connor Ludwig at Stanford University for building the photo-bleaching chamber used for histology analysis; Mr. Stefan Krabbe for building the custom lead helmet used in bone-marrow transplant experiment. Mr. Eric Martens for editing the manuscript. This work was supported by NIH grants to L.G. (1R01AG054214-01A1, U54NS100717, R01AG051390). The Gladstone Institutes received support from National Center for Research Resources Grant RR18928

## Methods

### Animals

All animal work was performed according to the approved guidelines from the University of California, San Francisco, Institutional Animal Care and Use Committee. Mice with *ad libitum* access to food and water were housed in a pathogen-free barrier facility with 12 hr light on/off cycle. C57BL/6 mice were used as wild-type control. Equal numbers of male and female mice were used for all experiments except for RNA-seq experiments, which used only female mice. Different mouse lines used in the study can be found in Supplementary table 2a.

### Drug administration

Diet containing 1200 mg/kg PLX5622 (Plexxikon Inc., Berkeley) was given to mice as the sole food source for 2-3 weeks to deplete microglia. Control diet with the same base formula but without the compound was given to the control group. Tamoxifen (Sigma-Aldrich, T5648) was prepared in corn-oil before use at 20 mg/mL. To efficiently induce Cre recombination, mice were given tamoxifen via intraperitoneal (IP) injection at 2 mg per day for 10 days. For sparse labeling in CX3CR1-CreERT2/Brainbow mice, tamoxifen was given via daily I.P. injection for 4 days. Mice receiving tamoxifen injections were housed for at least an additional 21 days after the last injection and before use. For labeling newborn cells, 5-Ethynyl-2’-deoxyuridine (EdU) (Santa Cruz, sc-284628) was prepared in sterile PBS at 20 mg/mL. EdU solution was warmed to 55 °C before use to dissolve any chemical precipitation. Mice were injected at 80 mg/kg via I.P. Diphtheria toxin (Sigma-Aldrich, D0564) was resuspended in sterile PBS at 20 ug/mL working solution for injection. DTX was delivered to animals via I.P. injection at 1 ug per day for 3 days.

### Bone marrow Transplantation

Bone marrow was isolated from the tibias and femurs of 3-6 months old ACTB-eGFP transgenic mice. The bone marrow was triturated using an 18-gauge needle and passed through a 70 μm nylon mesh cell strainer to make a single cell suspension. Erythrocytes were lysed with ACK lysis buffer (150 mM NH_4_Cl, 10 mM KHCO, and 0.1 mM Na_2_EDTA) and washed with PBS and suspended in PBS with 0.1% BSA at 3.5 × 10^7^ cells/mL. Recipient mice that were 4 to 6 months old were given two equal doses (2 × 600 rads, 3 hrs apart) of irradiation with a cesium source irradiator. To protect the brain from radiation, the head of the mouse was shielded by a lead plate during irradiation. Irradiated mice were reconstituted with 7 × 10^7^ cells of donor bone marrow via tail vein injection. Repopulation efficiency was determined by counting the percentage of GFP-positive myeloid population (CD45+CD11b+) by flow cytometry.

### Flow cytometry

Blood and spleen samples were collected in EDTA. Erythrocytes were lysed in FCK lysis buffer for 5 min and sample was pelleted at 500 g. Leukocytes were re-suspended in FACS buffer (PBS, 0.5% BSA, 5% FBA, 0.1% NaN3) and incubated for 15 min with anti-CD16/CD32 monoclonal antibodies (1:200, BD PharMingen) to block Fc receptors. Cells were then stained with APC-labeled anti-CD11b (1:100, Biolegend) and PE-labeled anti-CD45 (1:100, Biolegend) for 30 min on ice, followed by fixation with 2% PFA. Fluorescence intensity was measured using a FACS Calibur (BD Biosciences) flow cytometer. Data were analyzed in FlowJo (V10.0.7).

### Brain tissue harvest

Mice were anesthetized with avertin and transcardially perfused with PBS. Whole brains were drop-fixed in 4% paraformaldehyde prepared in PBS for 48 hr before switching to 30% sucrose for at least another 48 hr before cutting on a sliding microtome (Leica, SM2010R). Coronal sections of 30 μm thickness were collected. Sections were immersed in cyroprotectant and stored at −20 °C before use.

### Immunohistochemistry

One or two coronal sections per mouse were used for each staining. Free-floating sections were washed in PBS and then permeabilized in PBST buffer (0.5% Triton X-100 diluted in PBS) followed by blocking in 3% normal donkey serum (NDS) at room-temperature for 1 hr. Primary antibodies were diluted in PBST containing 3% NDS and incubated with tissues at 4 °C overnight. Primary antibodies used in the study can be found in Supplementary table 2b. Secondary antibodies were then prepared the same way as primary antibodies and incubated with tissue at room temperature for 1 hr. All secondary antibodies were obtained from Jackson ImmunoResearch. Secondary antibodies used in the study can be found in Supplementary table 2b. After secondary antibody staining, DAPI nuclear stain was conducted during the washing step as needed. Tissues were then mounted on glass slides for further processing or applied with anti-fade mounting media (Vector Laboratories, H-1000) for imaging. For detecting EdU+ cells in brain sections, Click-iT™ EdU imaging kits (ThermoFisher Scientific, C10337, C10339) were used following the manufacturer’s instructions before the immunofluorescence staining procedures. For terminal deoxynucleotidyl transferase-mediated dUTP nick end-labeling (TUNEL), the DeadEnd™ Fluorometric TUNEL System (Promega, G3250) was used with minor alterations from the manufacturer’s recommendations. We adapted a method developed by Deng *et al* [39]. In brief, after the immunofluorescence staining was finished, free-float brain sections were mounted on charged glass slide (Fisher, 12-550-15) and dried at 55 °C for 5 mins. Tissues were outlined with a hydrophobic pen (Vector Laboratories, H-4000). Mounted glass slides were then incubated in 0.5% Triton/PBS at 85°C for 20 min. After cooling to room-temperature, the equilibration buffer from the kit was applied directly to the slide for 5 min, followed by application of TUNEL reaction solution and incubation at 37 °C for 1 hr.

### Epifluorescence fluorescence microscopy

Regular epifluorescence images were acquired on a Keyence BZ-9000 inverted epifluorescence microscope equipped with an RGB and monochrome camera (Keyence, Osaka, Japan). Either the entire coronal hemi-brain slice or particular brain regions were scanned using 10x magnification and stitched in Keyence BZ-X Analyzer software (V1.3.0.3).

### Confocal fluorescence microscopy

Confocal microscopy was performed with a Zeiss LSM880 inverted scanning confocal microscope (Carl Zeiss Microscopy, Thornwood, NY) equipped with 2 PMT detectors, a high-sensitivity GaAsP detector, and a 32-GaAsP Airyscan super resolution detector, and run by Zeiss Zen imaging software. Confocal fluorescence images were acquired with 5-10 focal planes at 1-2 μm interval. Representative images are shown using Z-max intensity projections.

### Image analyses

All image analyses were performed in FIJI V1.50i [40]. For cell counting, tiff images were processed with adaptive threshold function to generate binary cell masks (https://sites.google.com/site/qingzongtseng/adaptivethreshold). Then the “Analyze Particles” function was used for cell counting. And cell density (per mm^2^) was calculated by normalizing cell number to the size of the analyzed area. For analyzing cells with multiple markers, the region of interest (ROI) was first generated from the base channel. Channels containing other makers of interest were thresholded to generate binary images. Each ROI generated from the base channel were then overplayed on to the binary images to calculate % area of the marker of interest. In general, >80% overlapping area is used to select ROIs that are positive for a second marker. For analyzing microglia 3D morphology, confocal z-stacks with 8 focal planes of 2 μm interval was used. Three 425.1×425.1μm^2^ connecting fields of view spanning the hippocampal CA1 to CA2 were captured and stitched in ZEN Blue software (Zeiss). The stitched tile images were analyzed in IMARIS software (V9.0.2, Bitplane). Microglia processes were analyzed using IMARIS filament function. Microglia soma size were measured using IMARIS surface function.

### Acute brain slice imaging

CX3CR1-GFP mice of 3-3.5 months old were treated for 2 weeks with a PLX-containing diet and then switched to regular diet to allow microglia repopulation for 6 days. Acute brain slices were prepared as previously described [41]. Mice were anesthetized with isoflurane and perfused with 20 mL of ice-cold, carbogen-saturated (95% O_2_, 5% CO_2_) NMDG-HEPES artificial cerebrospinal fluid (ACSF) solution containing: 93 mM NMDG; 2.5 mM KCl; 1.4 mM NaH_2_PO_4_; 30 mM NaHCO_3_; 20 mM HEPES; 25 mM D-glucose; 5 mM ascorbic acid; 2 mM thiourea; 3 mM sodium pyruvate; 12 mM N-acetyl-L-cysteine; 10 mM MgSO_4_; and 0.5 mM CaCl_2_. The ACSF was adjusted to pH 7.4 before use. All reagents were obtained from Sigma. Brains were washed in NMDG-HEPES. Coronal slices (200 μm thick) were prepared using a vibratome (Microm, Walldorf, Germany) at 4°C. For imaging, hemi-brain slices were transferred to a 35 mm glass bottom dish (MatTek) and secured under a slice anchor (Warner instruments). All imaging was done in non-NMDG ACSF solution containing (in mM): NaCl 92; KCl 2.5; NaH2PO4 1.4; NaHCO3 30; HEPES 20; D-glucose 25; Ascorbic acid 5; Thiourea 2; Sodium pyruvate 3; N-acetyl-L-cysteine 12; MgSO4 2; CaCl2 2; pH 7.4 (all from Sigma). Carbogen-saturated ACSF flowed continuously over the slice using a perfusion pump system (Cole-Parmer) at a rate of 3 mL/min. Images were captured with a Zeiss Z1 Observer inverted epifluorescence microscope (Carl Zeiss Microscopy, Thornwood, NY) with an ORCA-Flash 4.0 sCMOS camera (Hamamatsu Photonics, Shizuoka, Japan) and a Zeiss Axiocam MRm monochrome camera run by Zeiss Zen imaging software. The microscope is equipped with a motorized stage and temperature-controlled incubation system. Acquisition of cortical microglia was performed at a range of 50 to 100 μm from the surface of the slice to prevent capture of activated microglia, and z-stack images (1 μm step-size) were taken every 60 seconds for 15 to 30 minutes at 488 nm excitation and 510 nm emission wavelengths using a 40x objective. The motility of microglia processes was analyzed using the ImageJ plugin MTrackJ [42] to calculate the velocity of process retraction and extension per imaged cell.

### Microglia spatial pattern analyses

Hemi-brain slice from CX3CR1-CreERT2/Brainbow mice were stained with anti-GFP and anti-RFP antibody. The entire coronal section was scanned using a Keyence BZ-9000 inverted epifluorescence microscope. RFP+ or GFP+ cells were segmented and the XY coordinates were extracted from centeroids in FIJI. Nearest neighbor distances were calculated with ImageJ plugin NND (https://icme.hpc.msstate.edu/mediawiki/index.php/Nearest_Neighbor_Distances_Calculation_with_ImageJ). Spatial point pattern clustering analysis with Ripley’s K-function was performed as previously described [12]. The K-function is expressed as Equation 1, in which n is the number of total cells, *r* is the varying radius, *N_Pi_*(*r*) is the regional density of the *i*th cell at radius r, and *λ* is overall cell density. Theoretical complete spatial randomness (CSR) modeled by Poisson distribution in K(r) equals πr^2^. H-function, as shown in Equation 2, is transformed from the Ripley’s K-function for improved data visualization. In H-function, CSR equals zero. Clustering pattern is expressed as positive values whereas dispersion pattern is expressed as negative values.

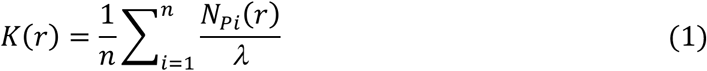

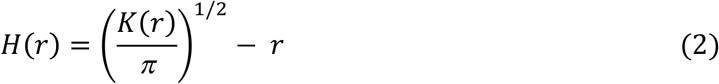

H(r) function was computed using Kest function in R package Spatstat [43]. H(r) values from each mouse were calculated separately. Average H(r) were then computed for each treatment group. Smoothing function using a moving average with interval of [r-15 μm, r+15 μm] was applied. Domain size was estimated from the corresponding radius at maximum of H(r) [23]. To define cluster boundaries, 2D kernel density estimates were calculated by using density function in Spatstat package. Spatial points of RFP+ or GFP+ cells were smoothed with 100 μm^2^ Gaussian kernel to generate density estimates. Regions containing the highest top 10% kernel density were chosen to generate cluster boundaries as this parameter best matches expected cell density. Overall size shift of the RFP+/GFP+ cluster overlapping region were quantified as percent of RFP and GFP overlapping area relative to the combined non-overlapping RFP and GFP area.

### Adult microglia isolation

Adult microglia isolation was performed using magnetic activated cell sorting (MACS) as previously described [41]. Briefly, mice were anesthetized with avertin and transcardially thoroughly perfused with PBS to remove circulating blood cells in the CNS. Each dissected brain was chilled on ice and then minced in enzymatic digestion buffer containing 3% Collagenase Type 3 (Worthington, LS004182) and 3 U/mL Dispase (Worthington, LS02104). Minced brain tissue was then incubated at 37 °C for 45 min. The enzymatic digestion was stopped with inactivation buffer containing 2.5 mM EDTA (Thermofisher, 15575020) and 1% fetal bovine srum (Invitrogen, 10082147). The digested brain tissue was then triturated in a serological pipette several times before passing through a 70 μm filter. The homogenate was then depleted with myelin using myelin removal beads (Miltenyi Biotec, 130-096-733) and a magnetic LD column (Miltenyi Biotec, 130-042-901). The elute was enriched for microglia with CD11b magnetic beads (Miltenyi Biotec, 130-049-601) and MS column (Miltenyi Biotec, 130-042-201). Microglial purity was determined by qPCR to access gene enrichment of microglial gene AIF1. Presence of the non-microglial fraction was assessed by measuring gene enrichment of NeuN and Aldh1l1.

### High-throughput RNA-sequencing

Female wild-type C57BL6 mice of 5-6 months old were used. To identify differential gene expression during microglia repopulation, mice were treated for 2 weeks with PLX-containing diet and then switched to regular diet allowing for microglia repopulation for various duration before being sacrificed for microglia isolation. These repopulation duration time points were set at 4 days (4D), 14 days (14D), and 1 month (1Mo). P4 microglia were isolated from 4-day-old postnatal mice as an immature microglia control. Microglia isolated from each single mouse were used as an individual sample for downstream steps. A total of 3 pups (P4), four untreated adult mice (ctrl) and 4 repopulated mice from each designed time point were used. Total RNA from fresh isolated microglia was extracted using Direct-zol RNA microprep kit (Zymo Research, R2061). cDNA library generation and RNA-seq service was performed by Novogene (Novogene Co., Ltd, Sacramento, California). RNA quality was examined by Bioanalyzer 2100 (Agilent Genomics). RNA samples with RIN value greater than 8 were used for cDNA library. Oligo(dT) beads were used to enrich for mRNA. After chemical fragmentation, a cDNA library was generated using NEBNext Ultra RNA Library Prep Kit for Illumina (New England Biolabs, E7530S). Quality of the cDNA library was assessed using Qubit assay for preliminary concentration, Bioanalyzer 2100 for insert size, and q-PCR for effective library size. QC passed cDNA library samples were then sequenced with the HiSeq 4000 system (Illumina) at PE150. On average, less than 0.01% error rate was detected and over 95% effective rate was observed in all sequencing results. Raw read ends containing low quality reading or adapter sequence were trimmed prior to downstream analysis.

### RNA-seq data analyses

RNA-seq read mapping was performed using the STAR program [44]. Gencode mouse genome GRCm38 was used as reference (release M16, 2017). On average, roughly 80% of reads were uniquely mapped to the reference genome. The read count table was generated with the RSEM program [45]. Differential gene expression was calculated with R package edgeR [46] and limma [47]. Genes which showed less than 1 count per million (CPM) in at least 3 samples were filtered out from further analysis. Normalization was performed with the weighted trimmed mean of M-values (TMM) method [48]. RNA-seq data were then transformed for linear model fitting with voom and lmFit functions inside limma package. Finally, empirical Bayes statistics were applied to correct variance of genes with low expression in the dataset. FDR was calculated by the Benjamini-Hochberg method [49]. DE genes were defined as two-fold change with false discovery rate (FDR) less than 0.05 in comparison to unperturbed adult microglia (Ctrl).

### Gene network and functional analyses

Gene network analyses were performed with gene set enrichment analysis (GESA) with molecular signatures database (MSigDB) [26, 27]. Network visualization was made in Cyotscape (version 3.6.0) with access to STRING database [50].

### Statistics

All experiments were performed with a minimum of at least 3 biological replicates. All data were averaged to individual animals. Mean values from each animal were used for computing statistical differences. Error bar in plots indicates standard error of the mean (SEM). Statistical analyses were performed in Graphpad prism 7.0c (Graphpad, San Diego, CA) and R (F Foundation for Statistical Computing, Vienna, Austria). Data visualization were achieved with R package ggplot2 [51]. Student’s t-test was used to compare two groups. One-way ANOVA was used to compare equal to or greater than three groups. Dunnett’s multiple comparisons test was used to compare difference with a specific group. Two-way ANOVA was used for groups with genotypes and treatment as factors. Sidak’s multiple comparisons test was used to compare statistical difference between genotypes. Statistical summary of p-values or FDR was indicated as p>0.05 (ns); p≤0.05 (*); p≤0.01 (**); p≤0.001 (***); p≤0.0001 (****).

## Author Contributions

Conceptualization, L.Z. and L.G.; Investigation, L.Z., G.K., M.C.R., L.K., S.H.C. and Y.L., Software, L.Z. F.D. and M.T.; Formal Analysis, L.Z. F.D. L.K. and M.T.; Writing – Original Draft, L.Z. and L.G.; Writing – Review & Editing, G.K., F.D., F.S., L.K. and L.G.; Funding Acquisition, L.G.; Resources, B.W., Y.S., F.S., D.L, Y.G., and C.W.; Supervision, L.G.

## Declaration of Interests

The authors declare no competing interests.

